# BDNF overexpression in the ventral hippocampus promotes antidepressant- and anxiolytic-like activity in serotonin transporter knockout rats

**DOI:** 10.1101/2020.07.01.181966

**Authors:** Danielle M. Diniz, Francesca Calabrese, Paola Brivio, Marco A. Riva, Joanes Grandjean, Judith R. Homberg

## Abstract

Brain-derived neurotrophic factor is one of the most studied proteins playing a pivotal role in neuroplasticity events and vulnerability and resilience to stress-related disorders. Most importantly, BDNF is decreased in depressive patients, and increased after antidepressant treatment. Additionally, BDNF was found to be reduced in a genetic subset of depression susceptible patients carrying the human polymorphism in the serotonin transporter promoter region (5-HTTLPR). The serotonin knockout rat (SERT^-/-^) is one of the animal models used to investigate the underlying molecular mechanisms behind the genetic susceptibility to depression in humans. SERT^-/-^ rats present decreased BDNF levels, especially BDNF exon IV, in the prefrontal cortex (PFC) and ventral hippocampus (vHIP), and display anxiety- and depression-like behavior. To investigate whether upregulating BDNF in the vHIP would meliorate the phenotype of SERT^-/-^ rats, we overexpressed BDNF locally into the rat brain by means of stereotaxic surgery and submitted the animals to behavioral challenges, including the sucrose consumption, the open field, and forced swim tests. Additionally, we measured hypothalamus-pituitary-adrenal (HPA)-axis reactivity. The results showed that lentivirus-induced BDNF IV overexpression in the vHIP of SERT^-/-^ rats promoted higher sucrose preference and sucrose intake, on the first day of the sucrose consumption test, indicative for decreased anhedonia-like behavior. Moreover, it decreased immobility time in the forced swim test, suggesting adaptive passive coping. Additionally, BDNF upregulation increased the time spent in the center of a novel environment, implying decreased novel-induced anxiety-like behavior. Finally, it promoted a stronger decrease in plasma corticosterone levels 60 minutes after restraint stress. In conclusion, modulation of BDNF IV levels in the vHIP of SERT^-/-^ rats led to a positive behavioral outcome placing BDNF upregulation in the vHIP as a potential candidate for the development new therapeutic approaches targeting the improvement of depressive symptoms.

## Introduction

An essential part of our daily performance has to do with our capability to adjust and respond to unforeseen events. Adaptation to unexpected environmental changes relies on our ability to develop resilience and display behavioral flexibility. This flexibility is a result of several mechanisms occurring in the brain that are usually called plasticity. Brain plasticity or neuroplasticity can be defined as the adjustments the nervous system undergoes in response to diverse stimuli to achieve reorganization at the structural, functional, or cellular connectivity level (Mateos-Aparicio and Rodríguez-Moreno, 2019). Considering this, it is expected that alterations in the brain’s ability to develop neuronal plasticity can lead to maladaptation and, consequently, to the development of neuropsychiatric disorders (Duman et al., 2000). Many molecular networks might be involved in neuroplasticity, including the serotonin and the brain-derived neurotrophic factor (BDNF) systems. Serotonin is a brain-wide distributed monoamine, and its decrease has been connected to the occurrence of depression (Coppen, 1967).

The levels of serotonin in the synaptic cleft are regulated by the serotonin reuptake transporter (SERT), and to restore serotonin levels in depressive subjects, drugs targeting SERT, known as selective serotonin-reuptake inhibitors (SSRI), are considered the first-line treatment for depressive disorders (Alenina and Klempin, 2015; Berger et al., 2009). While SSRIs very quickly generate a blockade of SERT in vitro, the actual clinic therapeutic effect may take several weeks (Deltheil et al., 2008). This delay suggests that the molecular mechanisms underlying depression might be more complex, requiring perhaps secondary changes in structural, functional, or cellular connectivity; thus, requiring neuronal plasticity (Duman et al., 2000, 1999).

BDNF, a neuropeptide from the neurotrophin family, plays a central role in neuroplasticity (Edelmann et al., 2014). Several studies confirmed that BDNF contributes to the brain plasticity through its positive influence in neurogenesis, cell survival, synapse formation and plasticity (Edelmann et al., 2014; Mattson et al., 2004; Park and Poo, 2013). BDNF gained attention in the study of mood disorders because data from post-mortem brain tissues or serum of depressive patients indicated low levels of BDNF, which were normalized following antidepressant treatment (Adachi, 2014; Sen et al., 2008). Given that the antidepressant therapeutic action occurs in a delay of weeks, and due to BDNF’s pivotal role in neuroplasticity, it has been associated with the mechanism of action of antidepressants (Duclot and Kabbaj, 2015; Duman and Monteggia, 2006; Monteggia et al., 2004). Therefore, the relation between the serotonin system and the BDNF system in connection with the antidepressant therapeutic effect seems to overlap.

The human SERT, encoded by the SLC6A4 gene, presents a functional polymorphism in the promoter region (5-HTTLPR) generating a short and a long allele variant of the serotonin transporter (Murphy et al., 2008). The short allelic variant induces a decrease in the SERT transcription in comparison to the long allelic variant (Lesch et al., 1996), and it is linked to increased risk for depression and suicidal behavior (Bleys et al., 2018; Caspi et al., 2003; Fanelli and Serretti, 2019; Homberg et al., 2014). Additionally, the short allele variant appears to be associated with insufficient response to SSRIs (Kroeze et al., 2012). Moreover, the short allelic variant has been related to decreased BDNF mRNA levels in white blood cells and serum from healthy 5-HTTLPR patients (Benedetti et al., 2017; Bhang et al., 2011; Molteni et al., 2010). Likewise, rodents lacking SERT (SERT^-/-^ animals) display a significant BDNF mRNA and protein downregulation, especially in the hippocampus and prefrontal cortex (Calabrese et al., 2015, 2013; Molteni et al., 2010). Additionally, SERT^-/-^ rats and mice present anxiety- and depression-related phenotypes (Kalueff et al., 2010; Olivier et al., 2008).

The hippocampus plays a crucial role in depression. The hippocampus modulates emotional processing, memory, learning, and controls glucocorticoid secretion by the hypothalamic-pituitary-adrenal axis (HPA-axis), making this area susceptible to the effects of stress (O’Leary et al., 2014). Stress and other negative stimuli can change hippocampal plasticity, increasing the risk of depression (Liu et al., 2017). In fact, depressive disorder is significantly associated with hippocampal atrophy (Elbejjani et al., 2015, 2014; Santos et al., 2018; Taylor et al., 2014). Furthermore, several lines of research have shown that BNDF and its high-affinity receptor (TrkB) are decreased in the hippocampus of post-mortem tissue from suicidal or depressed patients (Castrén et al., 2007; Castrén and Rantamäki, 2010; Dwivedi et al., 2003; Ray et al., 2011). Importantly, while impaired hippocampal neurogenesis can lead to depression (Jacobs et al., 2000), studies demonstrated that the upregulation of BDNF levels stimulated hippocampal neurogenesis (Quesseveur et al., 2013; Rossi et al., 2006).

In line with this, considering that BDNF levels are decreased in the ventral hippocampus of SERT^-/-^ rats (Calabrese et al., 2013; Molteni et al., 2010) we sought to investigate whether BDNF gene overexpression in the ventral hippocampus was able to restore BDNF levels and anxiety- and depression-like behavior in SERT^-/-^ rats. Although BDNF gene expression generates at least nine different transcripts (Aid et al., 2007), we targeted the overexpression of transcript IV because it is most downregulated in SERT^-/-^ rats (Molteni et al., 2010). Moreover, BDNF IV was selected due to its activity-dependent expression, association with depressive-like phenotypes (Sakata and Duke, 2014), and due to its higher contribution to total levels of BDNF protein in the hippocampus (Maynard et al., 2016). First, we aimed to analyze the temporal dynamics of BDNF IV overexpression one, two, and four weeks following lentivirus infusion in the ventral hippocampus (vHIP) of SERT^+/+^ rats. In this experiment, we measured BDNF mRNA levels of total BDNF, BDNF IV, and BDNF VI in the prelimbic and infralimbic cortices, and in the vHIP through RT-qPCR analysis. Thereafter, in a second experiment, following the viral infusions, SERT^-/-^ and SERT^+/+^ rats were submitted to behavioral testing. Behavior experiments included: (1) sucrose consumption test, in which we assessed the rats preference for sucrose over water and the total sucrose intake in grams; (2) forced swim test, where we scored for immobility, mobility and strong/high mobility behaviors; (3) open field, where we measured novelty-induced locomotor activity using Phenotyper® cages to evaluate distance moved, velocity, and time and frequency in the center; and (4) HPA-axis reactivity test, a test where the response of the HPA-axis upon acute restraint stress was checked through measurement of plasma corticosterone (CORT) levels.

## Material and Methods

### Animals

SERT^-/-^ rats (Slc6a41Hubr) were generated by N-ethyl-N-nitrosourea (ENU)-induced mutagenesis on a Wistar background (Smits et al., 2006) and outcrossed with commercially available Wistar^CrI:WI^ rats obtained from Charles River Laboratories (Horst, the Netherlands) for at least 15 generations. Ear punches were taken at the age of 21 days for genotyping, which was done by LGC (Hoddesdon, United Kingdom). SERT^+/+^ rats were used to check BDNF virus overexpression in naïve animals. For the behavioral experiments, male SERT^-/-^ and wild-type (SERT^+/+^) rats, weighing 350-400g at the beginning of the study were used (see experimental design in figure 1). All animals were housed in temperature-controlled rooms (21 °C) with standard 12/12-h day/night-cycle (lights on at 7:00 am) and food and water available ad libitum. 5-7 days before surgery, animals were socially housed in individually ventilated (IVC) cages for habituation. After surgery, animals were separately housed in the IVC cages until recovery. Animals were socially isolated during the sucrose consumption test; thereafter, the animals we socially housed again and kept under the same temperature and day/night-cycle throughout the entire experiment. All experiments were approved by the Committee for Animal Experiments of the Radboud University Nijmegen Medical Centre, Nijmegen, the Netherlands, and all efforts were made to reduce the number of animals used and their suffering.

**Figure 1.**
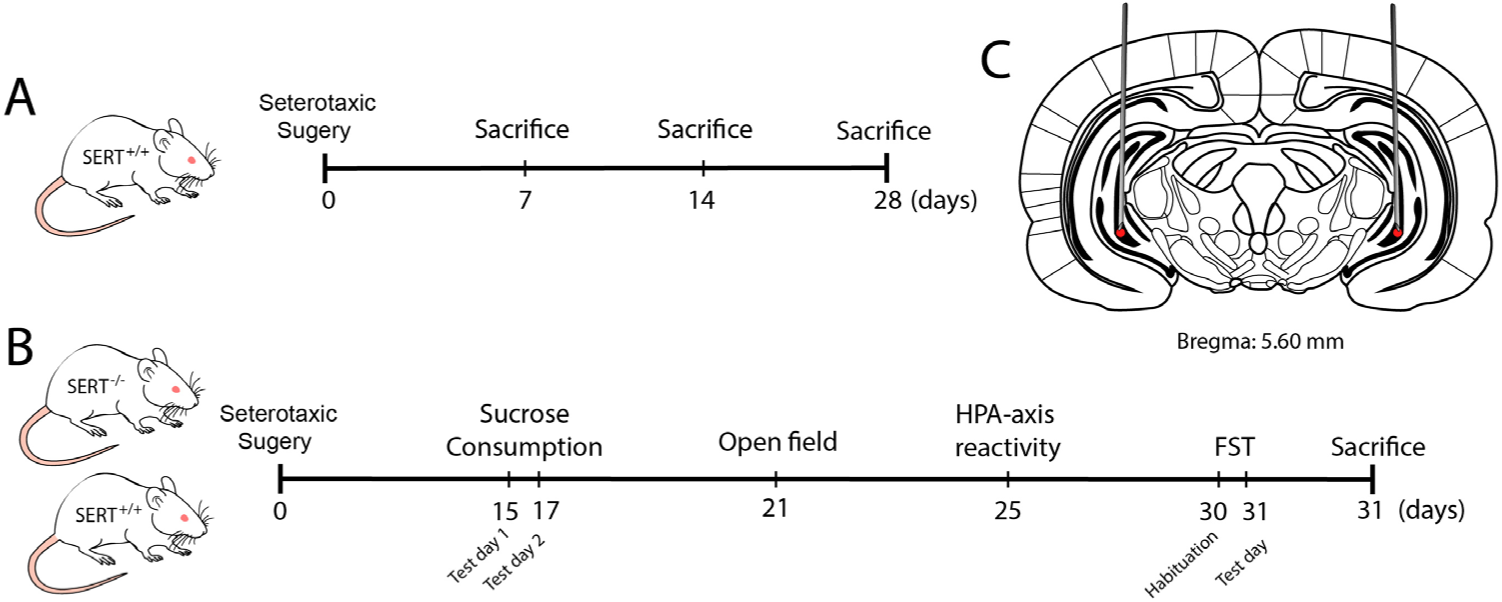
Schematic representation of the experimental design. A) Evaluation of BDNF overexpression in naïve SERT^+/+^ rats one, two, and four weeks following viral infusion. B) Behavioral tests: Viral infusion followed by behavioral tests including sucrose consumption test, HPA-axis reactivity, open field, and forced swim test (FST). C) Representation of the local o the infusion of either BDNF or control virus.

**Figure 2.**
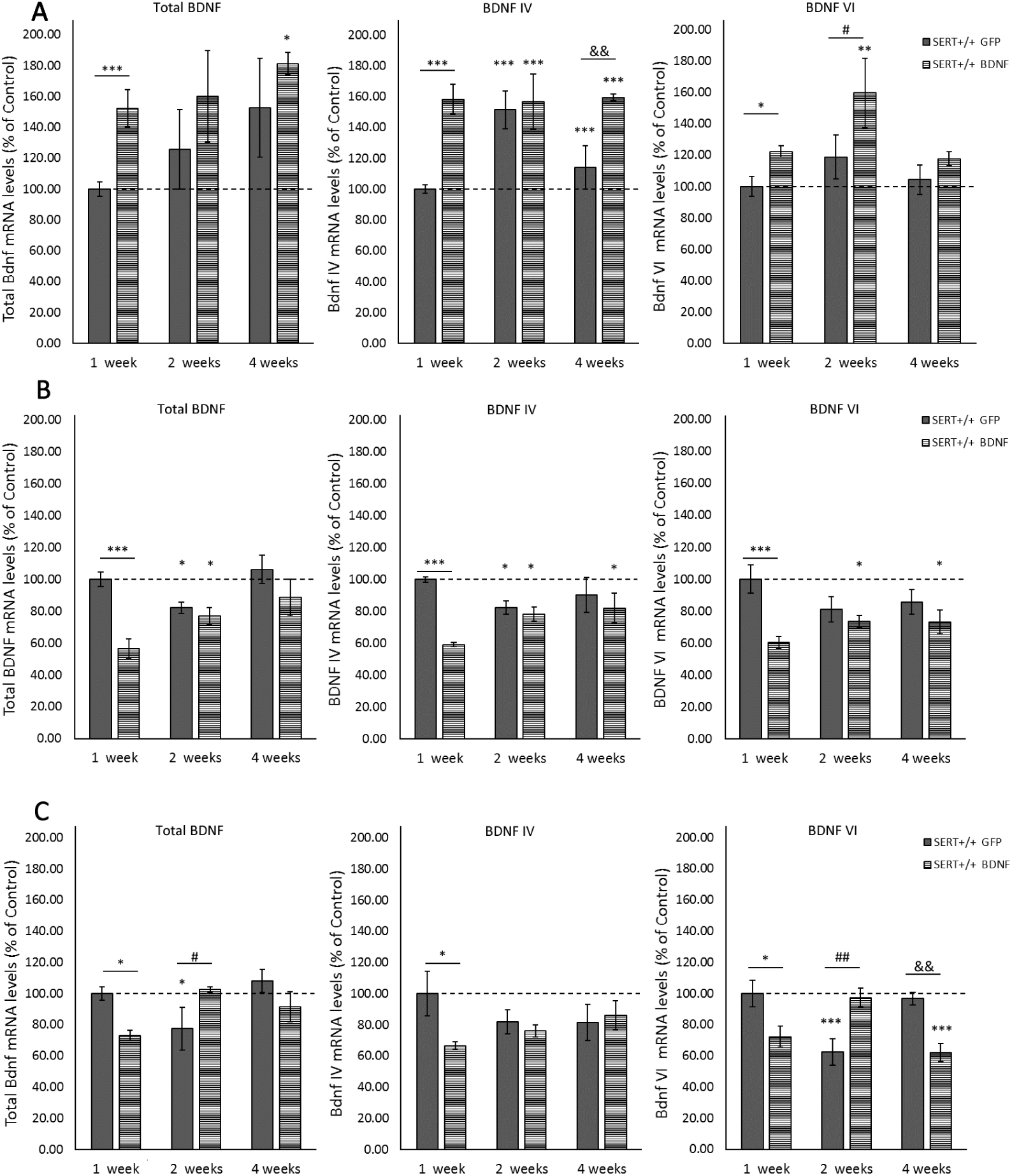
Modulation of total BDNF, BDNF IV, and BDNF IV transcripts expression in SERT +/+ animals infused with either GFP control or BDNF viral particles. BDNF mRNA levels were measured in the (A) ventral hippocampus, (B) infralimbic cortex, and (C) prelimbic cortex one, two, and four weeks after stereotaxic surgery. Data are expressed as fold change compared to the GFP-treated animals (set at 100%), and reflect mean ±SEM from 5-7 independent determinations. * = p < 0.05, ** = p < 0.01, and *** = p < 0.001 vs GFP-1 week; # = p < 0.05, ## = p < 0.01, and ### = p < 0.001 vs GFP-2 weeks; && = p < 0.01 vs GFP-4 weeks.

### Stereotaxic Surgery

Rats were anesthetized using isoflurane (5% induction, 2-3% maintenance). Lidocaine (10% m/v) was used for local anesthesia. Animals were fixed in a robot stereotaxic frame (StereoDrive, Neurostar, Germany). The coordinates for the site of the injection were theoretically determined based on the Paxinos & Watson (2007) rat brain atlas and checked through histological evaluation of 30 µm brain slices from dye-infused SERT^+/+^ rats. The total volume of 2μL of either BDNF lentivirus particles (transcript variant IV under CMV promoter, NM_001270633.1) or pLenti-C-mGFP control lentivirus particles, was bilaterally infused into the ventral hippocampus according to the following coordinates: AP -5.60 mm, ML ±5.0 mm, DV -7.6 mm. After surgery, animals were placed in IVC cages (Sealsafe Plus GR900 green line, Tecniplast, Italy) until sacrifice.

### RNA Preparation And Gene Expression Analysis By Quantitative Real-Time PC

Total RNA was isolated from the prelimbic and infralimbic region of the mPFC by single-step guanidinium isothiocyanate/phenol extraction using PureZol RNA isolation reagent (Bio-Rad Laboratories; Segrate, Italy), according to the manufacturer’s instructions, and then quantified by spectrophotometric analysis (NanoDrop(tm)1000, Thermo Scientific). Following total RNA extraction, an aliquot of each sample was treated with DNase to avoid DNA contamination. Then, the samples were processed for real-time PCR to assess total BDNF, BDNF isoform IV, and VI. The analyses were performed by TaqMan qRT–PCR instrument (CFX384 real-time system, Bio-Rad Laboratories S.r.l.) using the iScript one-step RT–PCR kit for probes (Bio-Rad Laboratories). Samples were run in 384-well formats in triplicates as multiplexed reactions with a normalizing internal control (36B4). Thermal cycling was initiated with incubation at 50°C for 10 min (RNA retrotranscription), and then at 95°C for 5 min (TaqMan polymerase activation). After this initial step, 39 cycles of PCR were performed. Each PCR cycle consisted of heating the samples at 95°C for 10 s to enable the melting process and then for 30 s at 60°C for the annealing and extension reactions. Data were analyzed with the comparative threshold cycle (ΔΔCt) method using 36B4 as a reference gene. Primers and probe for BDNF exon IV and VI were purchased from Life Technologies (BDNF exon IV: ID EF125679 and BDNF exon VI: ID EF125680). Primers and probe for total BDNF and 36B4 were purchased from Eurofins MWG-Operon. Their sequences are shown below:

- total BDNF: forward primer 5′-AAGTCTGCATTACATTCCTCGA-3′, reverse primer 5′-GTTTTCTGAAAGAGGGACAGTTTAT-3′, probe 5′-TGTGGTTTGTTGCCGTTGCCAAG-3′;

- 36B4: forward primer 5′-TTCCCACTGGCTGAAAAGGT-3′, reverse primer 5′-CGCAGCCGCAAATGC-3′, probe 5′-AAGGCCTTCCTGGCC GATCCATC-3′.

### Behavioral tests

#### Sucrose Consumption Test

After stereotaxic surgery, animals were housed individually and provided with two bottles of water for habituation. Before the test, side preference was checked for five days. The sucrose consumption test was adapted from Olivier et al. (2008) and consisted of two days of free-choice access to 24 hours sucrose versus water bottles with a water-only bottle choice in between the two days. In detail, on test day 1, one of the water bottles was replaced by sucrose 8% solution, and animals had free drinking access for a period of 24 hours. Next, animals received water in both bottles for 24 hours, ending with another 24 hours of free choice between water and sucrose 8% solution on test day 2. The position of the bottles was switched from sucrose consumption test day 1 to the test day 2 to prevent spatial bias. Daily, liquid intake and bodyweight were measured. The data are presented as the preference of sucrose above water (sucrose intake in ml divided by total intake X 100%) and the intake in grams of a 100% sucrose solution per kg bodyweight (intake in ml corrected for the voluminal weight of sucrose and recalculated toward a 100% solution divided by bodyweight in kg).

#### HPA-axis reactivity Test

The hypothalamic-pituitary-adrenal (HPA) axis reactivity was assessed through the measurement of corticosterone levels in the plasma. Usually, when rodents are submitted to stress, plasma concentrations of corticosterone (CORT) peak after 15 to 30 minutes and gradually decrease 60 to 90 minutes later to the pre-stress levels (de Kloet et al., 2005). Therefore, blood samples from tail cuts were collected in capillary blood collection tubes (Microvette® CB 300 Di-Kalium-EDTA, Sarstedt, Germany) 5 minutes before, and 15 and 60 minutes after 30 minutes of restraint stress. Rodent restrainers Broome-style were used for the restraint stress (554-BSRR, Bio-services, The Netherlands). Blood samples were centrifuged (3400 rpm for 15 min at 4 °C), and the plasma was stored at -80 °C until analysis. CORT levels were measured using a liquid chromatography-tandem mass spectrometry (LC-MS/MS).

#### Open field test

Novelty-induced locomotor activity was recorded by video recording in Phenotyper® cages (Noldus Information Technology, Wageningen, The Netherlands). The cages (45 cm × 45 cm × 45 cm) were made of transparent Perspex walls and a black floor. Each cage had a top unit containing a built-in digital infrared-sensitive video camera, infrared lighting sources, and hardware needed for video recording. To explore the novelty factor, animals were not exposed to this cage previously, and the cages were cleaned with 70% alcohol solution between trials to prevent transmission of olfactory cues. Spontaneous locomotor activity was monitored for 1 hour, and the following parameters were scored using Ethovision XT 11.5 (Noldus Information Technology, Wageningen, Netherlands): distance moved, velocity, frequency and time spent in the center of the cage (Manfré et al., 2017; Schipper et al., 2011).

#### Forced Swim test

The forced swimming test was performed as previously described (Porsolt et al., 1978). Briefly, rats were individually placed in cylindrical glass tanks (50 cm height, 20 cm diameter) filled to a height of 30 cm with 23±1°C water. The test consisted of two sessions. In the first session, animals were submitted to a habituation period of 15 minutes, then 24 hours later, to a second session of 5 min. The video recordings of the second session were used to automatically score the movements of the rats through a computerized system (Ethovision XT 10, Noldus, The Netherlands). Scored behaviors were ‘immobility’, which reflects no movement at all and/or minor movements necessary to keep the nose above the water; ‘mobility’, reflecting movement that corresponds to swimming activity; and ‘strong mobility’, reflecting ‘escape behavior’ (e.g., climbing against the walls and diving). Settings within Ethovision were adjusted based on manually recorded sessions (immobility/mobility threshold: 12; mobility/strong mobility threshold: 16.5 (Pawluski et al., 2016; Van den Hove et al., 2013).

### Statistical Analysis

The data were checked for outliers and normality (using the Shapiro–Wilk statistic), and extreme outliers were windsorized. Two-way analysis of variance (ANOVA) was computed for gene expression analysis, with time, genotype, and treatment as independent factors. The outcomes of sucrose consumption, open field, and forced swim tests, and post-behavioral gene expression were also analyzed through Two-way ANOVA considering genotype and treatment as fixed factors. Post-hoc Fisher’s Protected Least Significant Difference (PLSD) or independent sample t-tests were performed where applicable to compare individual group differences. All these statistical analyses were carried out using IBM^®^ SPSS^®^ statistics, version 23 (IBM software, USA). Regarding the HPA-axis reactivity test, a linear mixed model was implemented to account for repeated measurements, and multiple factor analysis using the LME4 package in R (3.5.1). Time, genotype, and treatment effects were modeled as a fixed effect, together with their pairwise double interactions, and their triple interactions. Subject intercepts were modeled as random effects. A likelihood-test ratio was used to assess fixed effect significance. Post-hoc tests were performed with the multicomp package, which accounts for multiple hypothesis testing. Significance was accepted at a *p*<0.05 threshold. Descriptive statistics are provided as mean +/-1 standard error of the mean (SEM).

## Results

### Lentivirus transfection leads to BDNF overexpression in ventral Hippocampus of SERT^+/+^ rats

Molecular analyses were performed to examine the feasibility of BDNF lentivirus overexpressing exon IV in the ventral hippocampus (vHIP) of naïve SERT^+/+^ rats. To check temporal dynamics, mRNA levels were evaluated one, two, and four weeks following BDNF or GFP lentivirus infusion. Due to the complexity of the BDNF gene and the distribution of its diverse transcripts variants across the brain (Aid et al., 2007), RT-qPCR was performed to measure total BDNF, BDNF IV and BDNF VI. Additionally, mRNA levels were measured not only in the vHIP but also in areas of high connectivity with the vHIP that are likewise involved in mood disorders, namely the prelimbic (PrL) and infralimbic (IL) cortex (Duman and Monteggia, 2006; Krishnan and Nestler, 2008; Schulz and Arora, 2015). The results from each brain area are detailed below.

#### Local BDNF IV lentivirus infusion leads to overall BDNF overexpression in the ventral hippocampus

Two-way ANOVA revealed significant main effects for the diverse BDNF transcripts analyzed. For example, a treatment main effect was observed for total BDNF (F_(1, 27)_ = 5.451, *p =* 0.027), BDNF IV (F_(1, 26)_ = 19.077, *p <* 0.001), and BDNF VI (F_(1, 27)_ = 8.389, *p =* 0.007). Moreover, an interaction between treatment and time was found for BDNF IV (F_(2, 26)_ = 4.533, *p =* 0.02). PLSD *post-hoc* analysis revealed that BDNF IV lentivirus infusion led to an expected upregulation of this transcript in the vHIP. BDNF IV levels in the site of the injection (vHIP) were significantly increased in SERT^+/+^ BDNF-treated animals compared to the control GFP-treated rats, one (*p <* 0.001), two (*p* < 0.001), and four weeks (*p <* 0.001) after surgery. Interestingly, two weeks after the viral infusion, we also identified a higher gene expression of transcript IV in control-treated SERT^+/+^ rats (*vs*. one-week GFP-treated animals, *p* < 0.001). Furthermore, as shown in figure A, pairwise comparisons also indicated an increase in total BNDF in SERT^+/+^ BDNF-treated rats one (*p* = 0.003) and for weeks after surgery (*p* = 0.032) in comparison to one-week GFP-treated animals. Additionally, BDNF VI levels were increased one (*vs* one-week SERT^+/+^ GFP, p = 0.02) and two weeks after the surgery compared to controls (*vs* one-week SERT^+/+^ GFP, *p* = 0.002; *vs* two weeks SERT^+/+^ GFP, *p* = 0.031). In conclusion, BDNF lentivirus infusion in the vHIP caused local overexpression of BDNF IV mRNA, which was stable for at least 4 weeks. Additionally, it led to total BDNF and BDNF VI upregulation in the vHIP.

#### BDNF overexpression in the vHIP leads to BDNF mRNA downregulation in the infralimbic cortex

A significant reduction in the analyzed BDNF transcripts in the IL was observed following the transfection of BDNF lentivirus into the vHIP. Two-way ANOVA was computed and results demonstrated main effects for treatment (F_(1, 28)_ = 16.756, *p <* 0.001), time (F_(2, 28)_ = 4.170, *p =* 0.026), and treatment *vs* time interaction (F_(2, 28)_ = 4.903, *p =* 0.015) for total BDNF levels. Additionally, treatment main effects were observed for BDNF IV (F_(1, 27)_ = 12.515, *p =* 0.001), and BDNF VI (F_(1, 29)_ = 12.514, *p =* 0.001). There was also a treatment *vs* time interaction (F_(2, 27)_ = 5.610, *p =* 0.009) for BDNF IV. Interestingly, as it can be seen in figure B, pairwise comparisons revealed that one week following the infusion in the vHIP, the levels of BDNF in the IL of BDNF-treated SERT^+/+^ rats dropped about 40% (*p* < 0.001) compared to control-treated animals at the same time point. Notably, both BDNF- and control-treated groups presented decreased levels of BDNF in the second week after the surgery (*p* = 0.05). This decrease was sustained for at least 4 weeks in BDNF-treated animals for BDNF IV (*p* = 0.038 *vs*. one-week SERT^+/+^ GFP-treated) and BDNF VI levels (*p* = 0.01 *vs*. one-week SERT^+/+^ GFP-treated), but total BDNF levels were back to normal four-weeks after the virus infusion. In conclusion, we observed that BDNF lentivirus infusion in the vHIP altered the BDNF levels in the infralimbic cortex, especially causing a significant decrease in BDNF mRNA expression one week after surgery.

#### BDNF overexpression in the vHIP alters BDNF expression in the prelimbic cortex

As in the IL, the levels of BDNF were decreased in the PrL of BDNF-treated SERT^+/+^ one week after the viral infusion in the vHIP. Specifically, main effects for the interaction between genotype and time point were found for total BDNF (F_(2, 28)_ = 6.044, *p =* 0.007) and BDNF VI (F_(2, 28)_ = 13.802, *p <* 0.001). *Post-hoc* analysis revealed that infusion of BDNF IV lentivirus particularly disturbed BDNF expression in a time- and transcript-dependent manner. For example, at the one-week time point, all the transcripts analyzed were decreased in the SERT^+/+^ rats treated with BDNF in comparison to controls (total BDNF *p* = 0.038, BDNF IV *p* = 0.02, and BDNF VI *p* = 0.014). However, as it is seen in figure C, two weeks following the surgery, while control rats presented decreased levels of total BDNF and BDNF VI (respectively *p* = 0.024 and *p* < 0.001 *vs*. one-week SERT^+/+^ GFP-treated), SERT^+/+^ rats treated with BDNF had these transcripts levels normalized to the control levels. Moreover, four weeks following the surgery, particularly the levels of BDNF VI were decreased in the BDNF-treated animals (*p <* 0.001 *vs*. one-week SERT^+/+^ GFP-treated, *p* = 0.003 *vs*. 4 weeks SERT^+/+^ GFP-treated). In short, BDNF overexpression in the vHIP caused decreased BDNF levels in the PrL one week after infusion; however, two and four weeks after, it especially altered the expression of BDNF VI resulting in both increased levels two weeks following surgery and decreased levels four weeks after surgery.

### Sucrose Consumption Test

Anhedonia is marked by a reduced interest in pleasurable events, and it is present in depression. This depression-like symptom can be identified in rodents through a decrease in sucrose consumption, which can be accessed through the sucrose consumption test. Animals were exposed to two days of free access to a sucrose 8% solution. The results of the preference for sucrose above the water and the sucrose intake in grams are described below.

#### Sucrose Preference: BDNF upregulation meliorates anhedonia-like behavior in SERT^-/-^ rats upon first exposure to the sucrose solution

Preference for sucrose was analyzed in two sessions. In the first day of test, two-way ANOVA indicated genotype as well treatment main effects (F_(1, 36)_ = 6.980, *p* = 0.012 and F(1, 36) = 4.854, *p* = 0.034, respectively). In detail, SERT^-/-^ rats treated with control virus presented a significant reduction in sucrose preference compared to both controls (*p* < 0.01) and BDNF-treated SERT^-/-^ rats (*p* = 0.006). Meanwhile, the BDNF-treated SERT^-/-^ rats displayed similar sucrose preference compared to control-treated SERT^+/+^ animals. On the second day, the BDNF treatment effect in SERT^-/-^ was not present anymore, and only a genotype effect was found (F_(1, 38)_ = 13.686, *p* < 0.001). Pairwise analysis indicated that both control- and BDNF-treated SERT^+/+^ had a higher preference for sucrose than both SERT^-/-^ groups (*p* < 0.05). In conclusion, as previously demonstrated, SERT^-/-^ rats displayed a lower preference for sucrose than SERT^+/+^ rats (Olivier et al., 2008); however, BDNF treatment improved the preference for sucrose in SERT^-/-^ rats at least in the first day of the test.

#### Sucrose intake: BDNF overexpression exclusively modulates sucrose intake on the first day of the sucrose consumption test

As done for the sucrose preference, sucrose intake was measured in two sessions. The results reveal that, on the first day of the test, BDNF lentivirus treatment altered the behavior of SERT^+/+^ and SERT^-/-^ regarding the consumption of sweet solution. This effect was lost on the second day. Two-away ANOVA showed a genotype (F_(1,38)_ = 29.313, *p <* 0.001) and a treatment effect (F_(1, 38)_ = 14.198, *p <* 0.001) on the first day of the test. In detail, both SERT^-/-^ and SERT^+/+^ rats treated with BDNF presented a significantly higher intake of sucrose than their respective GFP-treated controls (BDNF-*vs* GFP-SERT^-/-^, *p* = 0.017; and, BDNF-*vs* GFP-SERT^+/+^, *p* = 0.007). Moreover, compared to control SERT^+/+^ rats, only the GFP-treated SERT^-/-^ rats presented a lower intake of sucrose (*p* = 0.001). On the second day, just a genotype main effect was observed (F_(1, 38)_ = 23.280, *p <* 0.001). *Post-hoc* comparisons showed that BDNF treatment did not influence the intake in the second day and that – as previously described (Olivier et al., 2008) – SERT ^-/-^ rats consumed less sucrose than the SERT^+/+^ rats (*p* < 0.01). Taken together, the sucrose intake was influenced by the BDNF overexpression, with a positive effect on the anhedonia-like behavior of SERT^-/-^ rats and leading to an even higher sucrose intake in SERT^+/+^ rats. However, this effect was only present on the first day of the test. Upon the second exposition to the sucrose bottles, the SERT^-/-^ anhedonia-like phenotype returned.

### BDNF overexpression decreases immobility in the forced swim test in SERT^-/-^ rats

When rodents are exposed to an inescapable stressor such as in the forced swim test, their motivation to cope with stress can be quantified by the percentage of time spent on immobility (behavioral passivity) or performing a highly mobile (escape-like) behavior (Porsolt et al. 1977). Two-way ANOVA determined that for both extreme swimming modalities, immobility and high mobility, genotype main effects were found (F_(1,37)_ = 7.589, *p =* 0.009; and, F_(1,37)_ = 11.237, *p =* 0.002, respectively), while no main effects were found for the normal swimming (mobility) behavior scoring. Moreover, pairwise comparisons demonstrated that SERT^-/-^ rats were much more prompted to develop an escape-like behavior in comparison to SERT^+/+^ rats (*p* < 0.05). Furthermore, as shown in figure 3A, we observed that BDNF upregulation in the SERT^-/-^ rats resulted in a decreased immobility in comparison to both SERT^+/+^ groups (*vs*. SERT^+/+^ GFP, *p* = 0.025, and *vs*. SERT^+/+^ BDNF, *p* = 0.011). In conclusion, BDNF overexpression did not decrease the anxiety-like behavior in SERT^-/-^ rats expressed by increased time spent in the high mobility swimming modality; on the other hand, BDNF upregulation in the vHIP affected behavioral passivity, as expressed by a decrease in time spent on immobility in SERT^-/-^ rats.

**Figure 3.**
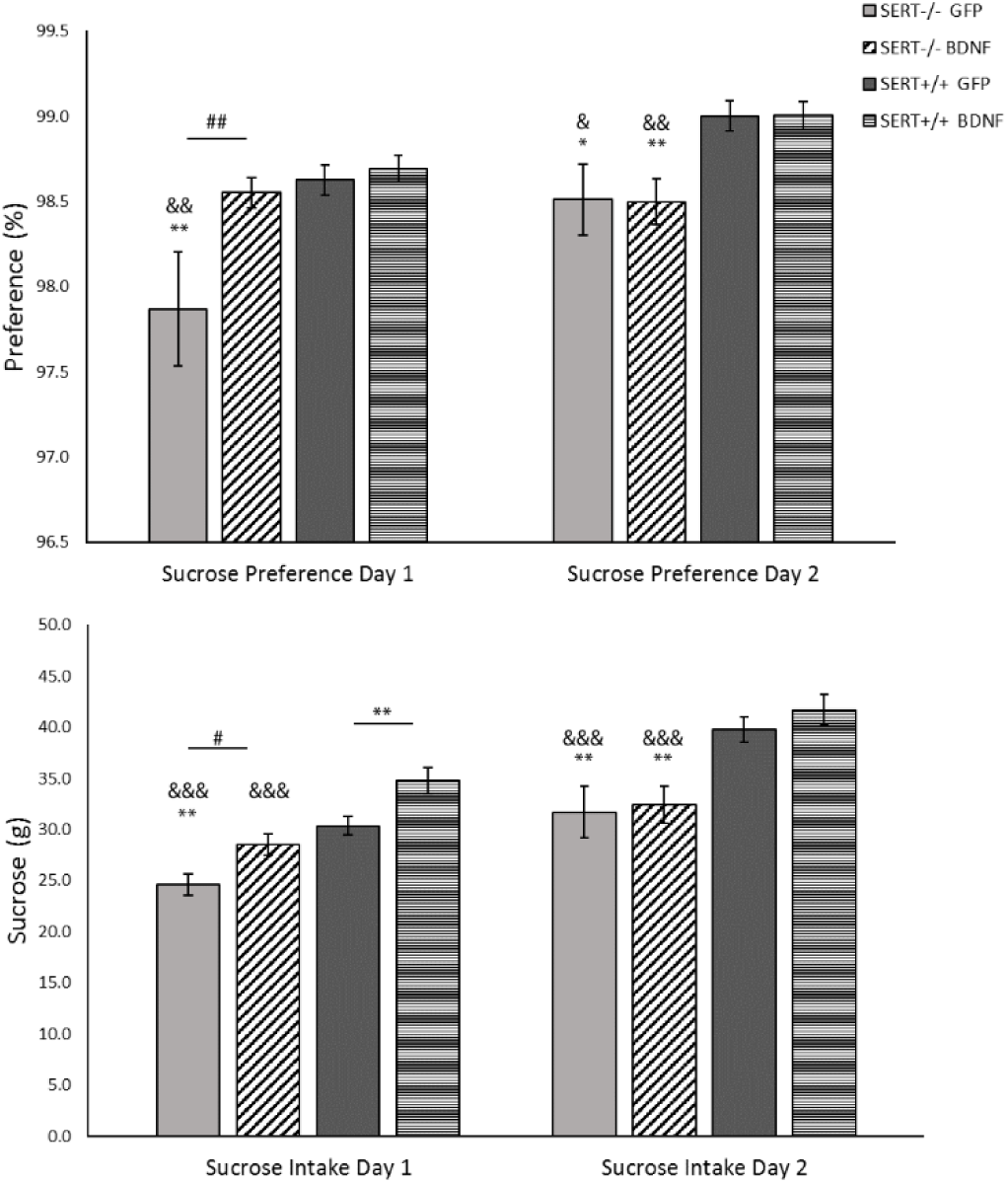
Sucrose consumption of 8% sucrose solution by SERT^-/-^ and SERT^+/+^ rats. Data are expressed as mean S.E.M. sucrose preference (sucrose intake/total fluid intake x 100%), and as mean S.E.M. total sucrose intake (g) per body weight (n = 10-12; * = p < 0.05 and ** = p < 0.01 vs SERT^+/+^-GFP; & = p < 0.05, && = p < 0.01, and &&& = p < 0.001 vs SERT^+/+^-BDNF; # = p < 0.05 vs SERT^-/-^GFP).

**Figure 4.**
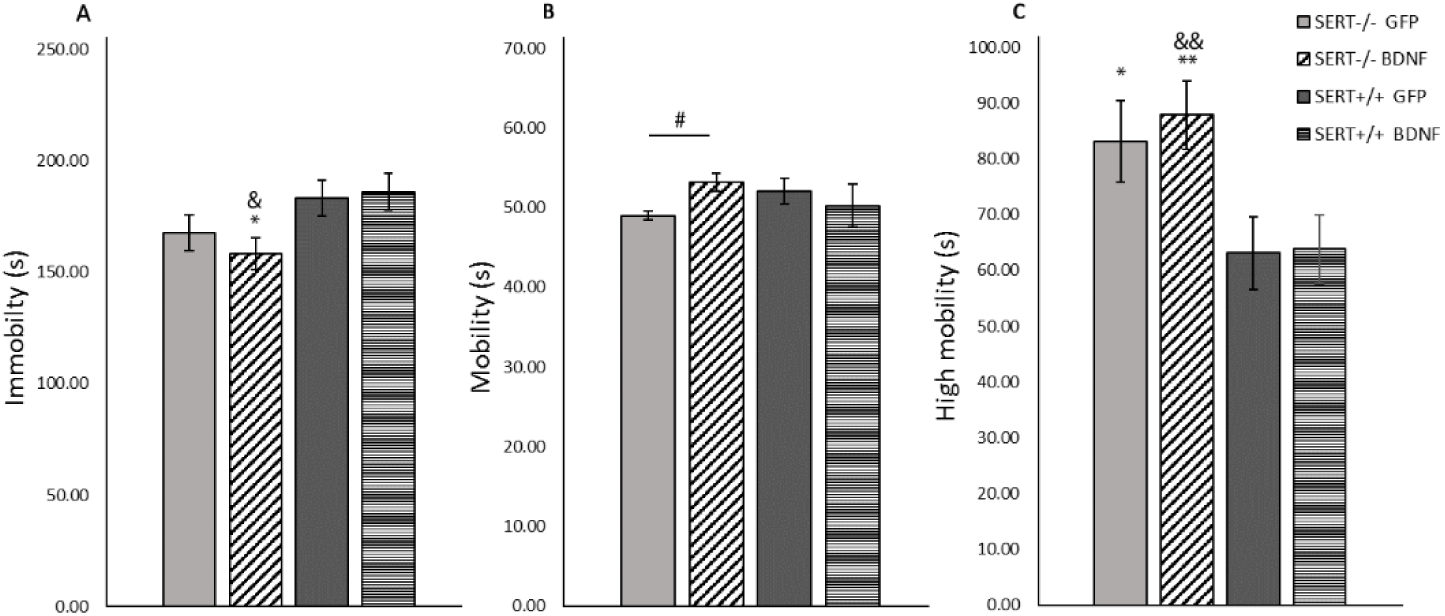
Mean (± SEM) measure of (A) immobility, (B) mobility, and (C) high mobility in the forced swim test (FST). n = 10–12 rats per group. * = p < 0.05, and ** = p < 0.01 vs SERT+/+ GFP; & = p < 0.05, and && = p < 0.01 vs SERT+/+ GFP; # = p < 0.05 vs SERT-/-GFP). Two-way ANOVA, Fisher LSD post-hoc test.

### BDNF upregulation decreases anxiety-like behavior in SERT^-/-^ rats in the open field test

Rodents may present a higher activity when they are introduced to a new environment (Menzaghi et al., 1994). A decrease in central locomotion (frequency and time spent in the central part of the arena), together with a general decrease in the locomotion (distance moved and velocity) can be interpreted as an anxiogenic-like behavior (Prut and Belzung, 2003). As previously reported, no differences between SERT^-/-^ and SERT^+/+^ rats were observed for the distance moved or velocity (Schipper et al., 2011). However, we did observe a genotype main effect for time spent in the center of the test-cage (F_(1, 37)_ = 6.557, *p* = 0.015). Further *post-hoc* analysis revealed that control-treated SERT^-/-^ rats spent less time in the center of the cage than control SERT^+/+^ rats (*p* = 0.007); meanwhile, BDNF injected SERT^-/-^ rats spent similar time in the center compared to the SERT^+/+^ rats. In conclusion, as reflected in figure 5, no alterations in the locomotor distance or speed were observed due to treatment or genotype. In contrast, the duration these animals spent in the center of the novel environment highlighted the behavioral differences between SERT^-/-^ and SERT^+/+^ rats. BDNF upregulation in SERT^-/-^ rats normalized the level of anxiety to that of SERT^+/+^ rats.

**Figure 5.**
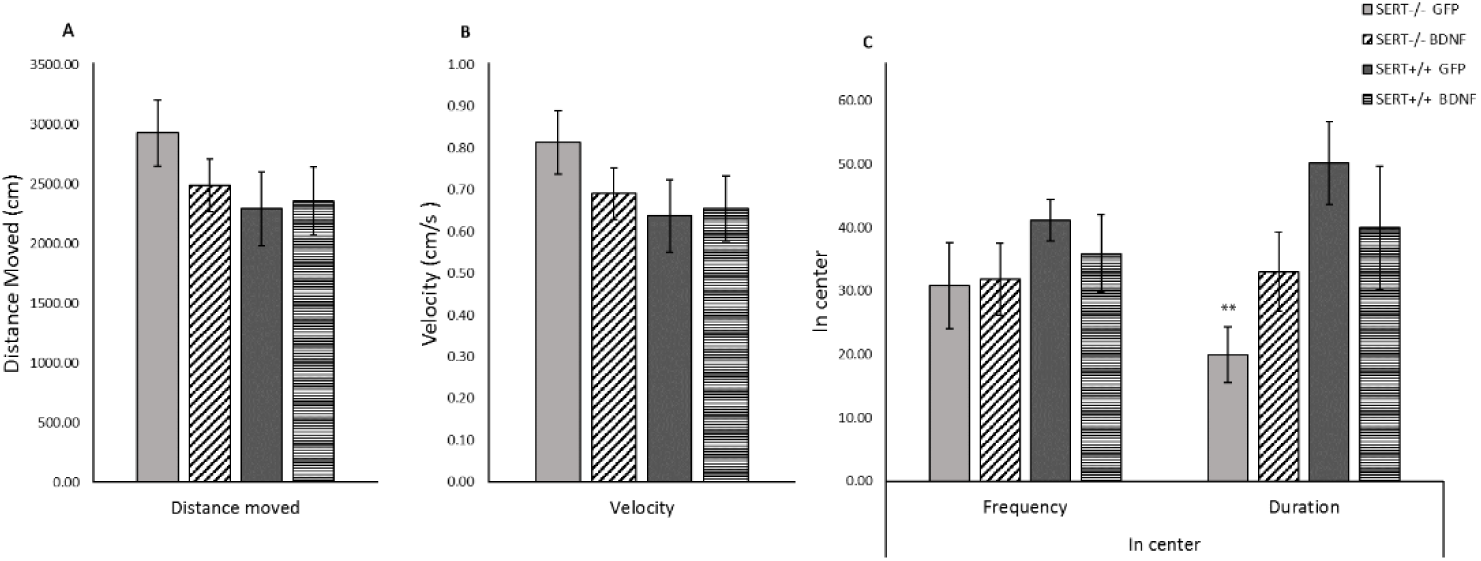
Novelty-induced locomotor activity expressed as mean (± SEM) (A) distance moved, (B) velocity, and (C) frequency and time spent in center. n = 10-12; ** = p < 0.01 vs SERT+/+ GFP). Two-way ANOVA, Fisher LSD post-hoc test.

**Figure 1.**
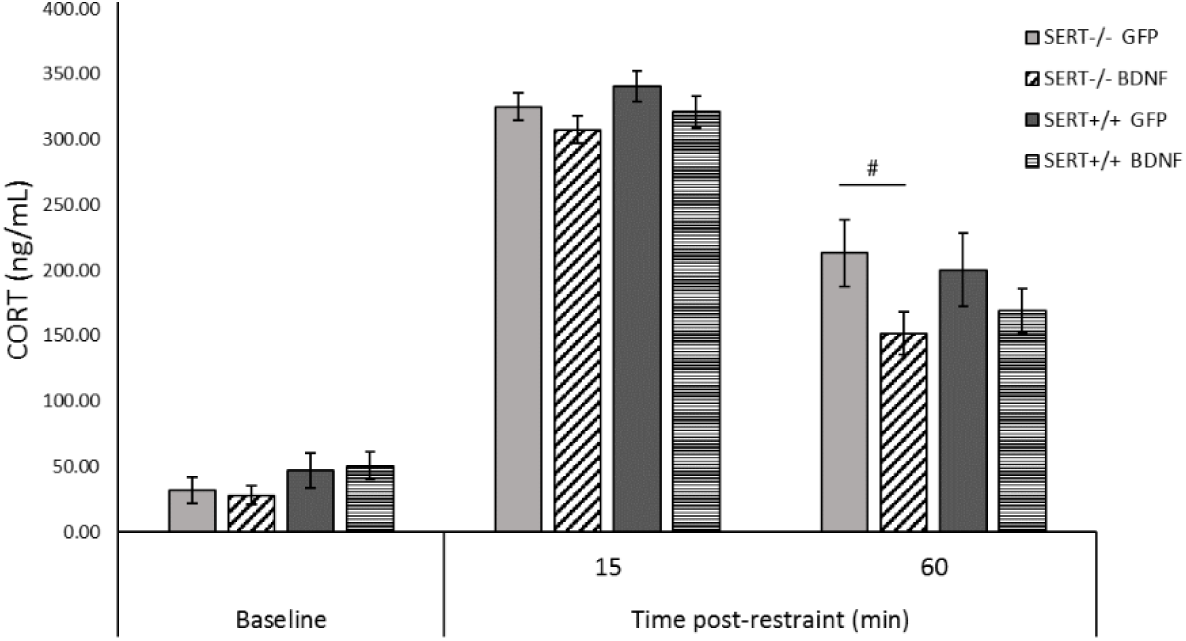
HPA-axis reactivity assessment. Corticosterone (CORT) levels are expressed mean ± (SEM) of plasma CORT levels 5 minutes before restraint stress (baseline), and 15 and 60 minutes post-restraint stress. n = 10-12; # = p < 0.05 vs SERT^-/-^ GFP. Linear mixed model.

### BDNF overexpression reduces CORT levels in SERT^-/-^ rats in the HPA-axis reactivity test

Acute restraint stress can activate the hypothalamic-pituitary-adrenal axis (HPA-axis), resulting in the release of CORT in rodents, which can be a measure of the (mal) functionality of the HPA-axis. The HPA-axis response to stress was evaluated 15 and 60 minutes after restraint stress. A linear mixed-effect analysis was conducted, and no interactions were found. However, pairwise comparisons indicated a significant decrease in CORT levels of BDNF-treated SERT^-/-^ rats 60 minutes after acute stress (*p* = 0.014), indicating that BDNF overexpression in the ventral hippocampus likely reduced the HPA reactivity in the SERT^-/-^ rats at this timepoint. No differences in the baseline CORT levels were found. Conclusively, while no basal or stress-induced differences in CORT levels were found between control-treated SERT^-/-^ and SERT^+/+^ rats, BDNF upregulation induced a decrease in the CORT levels in SERT^-/-^ rats versus control-treated SERT^-/-^ rats.

## Discussion

For many years, it has been hypothesized that the intricate system regulating the BDNF gene results in the generation of BDNF transcripts that create a spatial code for BDNF gene expression throughout the brain (Baj et al., 2011). Definitive conclusions regarding the role different transcripts play in BDNF signaling are still lacking, but it is of great interest giving that BDNF mRNA variants are differently (down)regulated in individuals suffering from neuropsychiatric disorders, including mood disorders. Considering this, we aimed to investigate the effects of BDNF IV overexpression in the ventral hippocampus of animals presenting both depressive-like behavior and BDNF downregulation. Here, we report that lentivirus BDNF IV infusion into the vHIP caused a significant overexpression of BDNF in the vHIP, followed by upregulation of other BDNF transcripts, such as total and BDNF VI levels. Interestingly, we observed that the upregulation of BDNF in the vHIP disturbed the BDNF gene expression in other brain areas. Particularly, we examined the PrL and IL and noticed an accentuated decrease in BDNF expression in these areas, especially in the first week following the BDNF viral infusion. Furthermore, we demonstrated that BDNF overexpression resulted in task- and genotype-dependent modifications in the phenotypic outcome from SERT^-/-^ and SERT^+/+^ rats. For instance, in the sucrose consumption test, BDNF treatment improved the preference for sucrose in SERT^-/-^ rats, and increased sucrose intake in SERT^-/-^ and SERT^+/+^ rats, but only in the first day of the test. Additionally, BDNF-infused SERT^-/-^ rats presented decreased immobility in the forced swim test, and lower anxiety levels in the novelty-induced locomotor activity test compared to controls. Finally, BDNF upregulation induced a decrease in HPA-axis activity, reflected by the decreased CORT levels in SERT^-/-^ rats.

Precise time-, brain region-, and stimuli-dependent BDNF production might be mediated by the expression of multiple BDNF mRNA variants (Maynard et al., 2016). At least 9 promoters are involved in the regulation of BDNF (Aid et al., 2007); among them, promoter IV is an activity-dependent gene, leading to the expression of BDNF upon neuronal activity (Sakata et al., 2010). Interestingly, molecular analysis of the BDNF IV overexpression in this study revealed exciting features regarding the BDNF IV effect over other BDNF transcripts, and regarding the vHIP connection with frontal areas of the rat brain. In short, one week after the viral infusion into the vHIP, we demonstrated that total BDNF, BDNF IV, and BDNF VI were overexpressed in the vHIP and downregulated in the PrL and IL of BDNF-infused SERT^+/+^ rats. In the IL, the decreased BDNF levels of variants IV and VI were stable for at least four weeks, showing a continuous influence from the vHIP BDNF upregulation. On the other hand, in the PrL, we noticed a time-dependent control specific to total and BDNF VI. Two weeks after BDNF infusion, PrL levels of total BDNF and BDNF VI were higher in BDNF-infused animals, but in the fourth week, BDNF VI levels were again downregulated. The medial prefrontal cortex (mPFC) receives direct projections from the ventral hippocampus (Verwer et al., 1997). Studies in cultured hippocampal neurons revealed activity-induced BDNF dendritic release (Adachi, 2014; Edelmann et al., 2014; Matsuda et al., 2009). Altogether, these data indicate that, likely, BDNF upregulation in the vHIP caused negative feedback in the BDNF transcription of downstream targets such as the PrL and IL. Importantly, just as the vHIP-mPFC synchrony was shown to be modulated by negative emotions like anxiety (Adhikari et al., 2010), the vHIP to mPFC transcriptional control, might likewise induce effects on cognitive outcome. Furthermore, the different effects in PrL and IL BDNF gene expression might specifically modulate cognitive (mal)functions, considering that these two mPFC subdivisions have different projections sites (Vertes, 2004; Vidal-Gonzalez et al., 2006).

Anhedonia is a core symptom in mood disorders (Morris et al., 2017). This negative emotional state has been previously demonstrated in SERT^-/-^ rats (Olivier et al., 2008), and was replicated in this study. In our experimental conditions, control-treated SERT^-/-^ rats displayed anhedonia-like behavior when compared to controls SERT^+/+^ rats. This anhedonia-like behavior was suppressed in the SERT^-/-^ rats following BDNF upregulation. However, the enhancement in sucrose preference and sucrose intake was limited to the first day of the test, and it was lost on the second day of the test. Therefore, the upregulation of BDNF IV in the vHIP likely ameliorated the anhedonia-like symptoms in SERT^-/-^ rats only when the novelty factor was present. After habituation to the novel sweet taste, the anhedonic phenotype was prevalent in SERT^-/-^ animals in comparison to SERT^+/+^ rats. Importantly, in a previous experiment (Diniz et al. unpublished data), we overexpressed BDNF in the PrL, and we observed exactly the opposite effect on sucrose preference. In short, SERT^-/-^ rats infused with BDNF IV into the PrL preferred less sucrose than GFP-infused SERT^-/-^ rats only on the first day of the sucrose consumption test. On the second day, the BDNF-treated animals showed higher sucrose preference than GFP-treated SERT^-/-^ rats. Therefore, BDNF upregulation differently modulated the response to a novel taste depending on the targeted brain region, causing both decreased sucrose preference in the PrL-infused animals, and increased sucrose over water preference in vHIP-infused animals.

Another core endophenotype in mood disorders is behavioral despair (Kalueff et al., 2010). In rodents, this symptom can be evaluated through the forced swim test (Porsolt et al., 1978). We demonstrated that, in our experimental conditions, we did not observe differences in immobility scores when comparing control-treated SERT^-/-^ to SERT^+/+^ rats. This result is at odds with previous studies using SERT^-/-^ rats and mice, in which immobility was increased in knockout animals (Lira et al., 2003; Olivier et al., 2008). However, under the same experimental conditions, SERT^-/-^ rats submitted to stereotaxic surgery targeting GFP virus infusion in the PrL also did not present increased immobility behavior in the forced swim test (Diniz et al. unpublished observations). Interestingly, BDNF upregulation in the vHIP decreased immobility time in SERT^-/-^ rats in comparison to GFP-treated SERT^-/-^ animals, suggesting that BDNF overexpression contributed to an adaptive learned response (Anyan and Amir, 2018). In line with this finding, Karpova et al. (2009) demonstrated that an increase in hippocampal BDNF IV caused by postnatal SSRI exposure was associated with decreased immobility in the forced swim test. Further, we demonstrated that SERT^-/-^ rats, independently of the treatment, displayed higher escape-behavior than SERT^+/+^ rats. Increased strong mobility in the forced swim test might indicate enhanced anxiety-like behavior displayed by SERT^-/-^ rats. Anxiety-like behavior in SERT^-/-^ rats might be due to changes in the functioning of the GABAergic system in SERT^-/-^ rats (Calabrese et al., 2013; Guidotti et al., 2012; Luoni et al., 2013; Miceli et al., 2017; Schipper et al., 2019), causing GABA downregulation and consequently, increase anxiety-like behavior (Lydiard, 2003; Millan, 2003).

In line with this observation, we identified that in the novelty-induced locomotor activity test, SERT^-/-^ rats spent less time in the center of the test-cage than SERT^+/+^ rats, indicating likewise, increased anxiety-like behavior (Olivier et al., 2008). Interestingly, SERT^-/-^ rats receiving BDNF virus as treatment did not differ from SERT^+/+^ controls, indirectly suggesting that BDNF upregulation likely decreased the SERT^-/-^ rats’ anxiety-like behavior in this behavior paradigm. This BDNF anxiolytic-like action modulated by exposition to novelty is in agreement with the results from the sucrose consumption test, in which SERT^-/-^ rats seems to present enhanced negative emotionality in connection to the novel taste and this behavior was reduced by BDNF IV overexpression in the vHIP. Additionally, in our experimental conditions, we replicated the previous observation that no alterations in the locomotor distance or speed were observed in SERT^-/-^ rats in comparison to SERT^+/+^ rats (Homberg et al. 2008; Schipper et al. 2011). This also indicates that the decreased immobility and increased high mobility observed in SERT^-/-^ rats in the forced swim test was not due to increased locomotor activity.

Adaptive responses to stressors require behavioral flexibility that depends on a plastic brain. Considering this, we also investigated the effects of hippocampal upregulation of BDNF IV on HPA-axis activity through the measurement of corticosterone (CORT) levels following acute stress. We demonstrated that 60 minutes after the acute stress, BDNF upregulation in the vHIP caused a significant decrease in CORT levels in SERT^-/-^ rats compared to GFP-treated SERT^-/-^ rats, indicating that BDNF overexpression affected the decreasing phase in CORT levels (de Kloet et al., 2005) by enhancing the negative feedback over the HPA-axis. Regarding the peak in CORT levels, at 15 minutes following the acute restraint stress, we observed that CORT levels were not affected by BDNF upregulation or genotype. Likewise, basal CORT levels were comparable in SERT^-/-^ and SERT^+/+^ rats, denoting a lack of BDNF modulation in baseline levels. In agreement with these results, studies in SERT^-/-^ mice also have shown that plasma basal CORT levels (Chen et al., 2012; Li et al., 1999; Tjurmina et al., 2002) or stress-response CORT levels (Jansen et al., 2010) were not altered in comparison to SERT^+/+^ mice. However, in contrast with our results, van der Doelen et al. (2014) have shown that non-stressed SERT^-/-^ rats presented increased baseline levels of CORT compared to SERT^+/+^ rats, whereas when submitted to early life stress, SERT^-/-^ rats present lower levels of CORT than early-life-stressed SERT^+/+^ rats. The data found in animals do not always match those from in human studies; however, basal HPA hyperactivity, and consequently hypercortisolemia is commonly detected in depressive patients (Nestler et al., 2002; Pariante and Lightman, 2008). In humans, the short allele generated by the single polymorphism in the serotonin transporter gene (5-HTTLPR) is associated with higher levels of waking cortisol (CORT) and increased CORT response to stress (Gotlib et al., 2008; Klein Gunnewiek et al., 2018; O’Hara et al., 2007). Therefore, the negative emotionality and neurodevelopmental changes presented by the SERT^-/-^ rodents are comparable to the effects of the short allele in the human 5-HTTLPR (Homberg et al., 2014; Kalueff et al., 2010). In our experimental conditions, the SERT^-/-^ rats were not stressed in their early life stage; however, they underwent surgery stress, isolation stress (in the sucrose consumption test), and were exposed to a novel environment in the locomotor activity test. Therefore, although animals were given time to recover, it is possible that the sequence of behavioral tests shaped an adaptive CORT response to stress (Belay et al., 2011) as well as determined the differential CORT basal levels (van der Doelen et al., 2014). Moreover, while BDNF overexpression in the vHIP did not alter baseline CORT levels in SERT^+/+^ rats, our previous data on BDNF overexpression in the prelimbic cortex of WT rats indicated significant increased basal CORT levels in these animals. Besides, in contrast to the BDNF overexpression in the vHIP, upregulation in the prelimbic cortex modulated the stress CORT levels 15 minutes following the acute restraint stress, causing increased CORT levels in SERT^-/-^ rats and decreased CORT levels in WT animals. Altogether, these data suggest that BDNF affects the HPA-axis activity in a brain region and phase-dependent manner; also, it differently affected the animals depending on their genotype. Further studies are necessary to understand the association between the serotonin transporter gene and the HPA-axis reactivity in stress and non-stress contexts. Especially, the role of BDNF in key brain areas controlling the HPA-axis response, such as is the case of the mPFC and hippocampus, in healthy and disease will be essential to develop novel treatments to provide improved care to patients suffering from mood disorders.

Depression is a complex neuropsychiatric disorder comprising multiple neural circuit processes (Nestler et al., 2002). Animal models such as the SERT^-/-^ rat comprise a useful tool to study targeted circuits that are found in both rodents and humans. Although complete function loss of the human SERT is a very rare condition, behavioral and biomolecular changes in SERT^-/-^ are comparable to short allele carriers of the human 5-HTTLPR (Kalueff et al., 2010). The SERT knockout rats are characterized by a complete lack of SERT and increased extracellular serotonin levels (Homberg et al. 2007), yet show anxious and depressive-like phenotypes. In agreement with the neurotrophic hypothesis of depression, these animals present, under basal conditions, downregulation of BDNF mRNA (especially exon IV) and protein levels in the ventral hippocampus and prefrontal cortex (Calabrese et al., 2013; Molteni et al., 2010). Modulation of BDNF IV levels in the ventral hippocampus and not in the prefrontal cortex showed to have positive effect in mood disorder-related symptoms. Selective upregulation of a specific BDNF exon can lead to specific neuronal network reinforcement (Baj et al., 2011). The current study demonstrated that defining a target brain region might be critical for mediating therapeutic interventions targeting BDNF overexpression. Further characterization of the effects BDNF overexpression might exert over other brain areas targeted by the vHIP, such as the PFC and nucleus accumbens (NAc), is still needed. In addition, given that BDNF did not decrease escape behavior in the forced swim test, indicating anxiety-like behavior, it is still unclear which circuits might be involved in the forced swim test escape-behavior response. Moreover, it is uncertain whether BDNF overexpression in the vHIP restored the decreased GABAergic inhibitory regulation in SERT^-/-^ rats (Calabrese et al., 2013; Luoni et al., 2013; Miceli et al., 2017; Schipper et al., 2019). Nonetheless, despite necessary further investigations, we hypothesize that BDNF IV upregulation in the vHIP might be a good candidate for modulation in the treatment of mood disorders. Performing stereotaxic surgery in humans seems not to be a feasible treatment for mood disorders in humans; however, advances in pharmaceutical technology have shown that the delivery of drugs and genes into the brain through peripheral bloodstream route is possible (Saraiva et al., 2016). Therefore, the additional studies to understand the BDNF dynamics in the vHIP of SERT^-/-^ rats might enhance the possibilities to the development of specific gene target delivery, allowing a future novel, rational treatment to bring individualized and improved care for patients with mood disorders.

## Funding

This work was supported by the Science Without Borders scholarship program from CNPq - Conselho Nacional de Desenvolvimento Científico e Tecnológico, of the Ministry of Science, Technology and Innovation of Brazil, grant #200355/2015-5, awarded to DMD.

## Notes

### Competing Interest Statement

The authors have declared no competing interest.

## References

1. Adachi, N., 2014. New insight in expression, transport, and secretion of brain-derived neurotrophic factor: Implications in brain-related diseases. World J. Biol. Chem. 5, 409. https://doi.org/10.4331/wjbc.v5.i4.409

2. Adhikari, A., Topiwala, M.A., Gordon, J.A., 2010. Synchronized activity between the ventral hippocampus and the medial prefrontal cortex during anxiety. Neuron 65, 257–69. https://doi.org/10.1016/j.neuron.2009.12.002

3. Aid, T., Kazantseva, A., Piirsoo, M., Palm, K., Timmusk, T., 2007. Mouse and ratBDNF gene structure and expression revisited. J. Neurosci. Res. 85, 525–535. https://doi.org/10.1002/jnr.21139

4. Alenina, N., Klempin, F., 2015. The role of serotonin in adult hippocampal neurogenesis. Behav. Brain Res. 277, 49–57. https://doi.org/10.1016/j.bbr.2014.07.038

5. Anyan, J., Amir, S., 2018. Too Depressed to Swim or Too Afraid to Stop? A Reinterpretation of the Forced Swim Test as a Measure of Anxiety-Like Behavior. Neuropsychopharmacology 43, 931–933. https://doi.org/10.1038/npp.2017.260

6. Baj, G., Leone, E., Chao, M. V., Tongiorgi, E., 2011. Spatial segregation of BDNF transcripts enables BDNF to differentially shape distinct dendritic compartments. Proc. Natl. Acad. Sci. 108, 16813–16818. https://doi.org/10.1073/pnas.1014168108

7. Belay, H., Burton, C.L., Lovic, V., Meaney, M.J., Sokolowski, M., Fleming, A.S., 2011. Early Adversity and Serotonin Transporter Genotype Interact With Hippocampal Glucocorticoid Receptor mRNA Expression, Corticosterone, and Behavior in Adult Male Rats. Behav. Neurosci. https://doi.org/10.1037/a0022891

8. Benedetti, F., Ambrée, O., Locatelli, C., Lorenzi, C., Poletti, S., Colombo, C., Arolt, V., 2017. The effect of childhood trauma on serum BDNF in bipolar depression is modulated by the serotonin promoter genotype. Neurosci. Lett. 656, 177–181. https://doi.org/10.1016/j.neulet.2017.07.043

9. Berger, M., Gray, J.A., Roth, B.L., 2009. The Expanded Biology of Serotonin. Annu. Rev. Med. 60, 355–366. https://doi.org/10.1146/annurev.med.60.042307.110802

10. Bhang, S., Ahn, J.-H., Choi, S.-W., 2011. Brain-derived neurotrophic factor and serotonin transporter gene-linked promoter region genes alter serum levels of brain-derived neurotrophic factor in humans. J. Affect. Disord. 128, 299–304. https://doi.org/10.1016/J.JAD.2010.07.008

11. Bleys, D., Luyten, P., Soenens, B., Claes, S., 2018. Gene-environment interactions between stress and 5-HTTLPR in depression: A meta-analytic update. J. Affect. Disord. 226, 339–345. https://doi.org/10.1016/J.JAD.2017.09.050

12. Calabrese, F., Guidotti, G., Middelman, A., Racagni, G., Homberg, J., Riva, M.A., 2013. Lack of Serotonin Transporter Alters BDNF Expression in the Rat Brain During Early Postnatal Development. Mol. Neurobiol. 48, 244–256. https://doi.org/10.1007/s12035-013-8449-z

13. Calabrese, F., van der Doelen, R.H.A.A., Guidotti, G., Racagni, G., Kozicz, T., Homberg, J.R., Riva, M.A., 2015. Exposure to early life stress regulates Bdnf expression in SERT mutant rats in an anatomically selective fashion. J. Neurochem. 132, 146–154. https://doi.org/10.1111/jnc.12846

14. Caspi, A., Sugden, K., Moffitt, T.E., Taylor, A., Craig, I.W., Harrington, H.L., McClay, J., Mill, J., Martin, J., Braithwaite, A., Poulton, R., 2003. Influence of life stress on depression: Moderation by a polymorphism in the 5-HTT gene. Science (80-.). 301, 386–389. https://doi.org/10.1126/science.1083968

15. Castrén, E., Rantamäki, T., 2010. The role of BDNF and its receptors in depression and antidepressant drug action: Reactivation of developmental plasticity. Dev. Neurobiol. 70, 289–297. https://doi.org/10.1002/dneu.20758

16. Castrén, E., Võikar, V., Rantamäki, T., 2007. Role of neurotrophic factors in depression, Current Opinion in Pharmacology. Elsevier. https://doi.org/10.1016/j.coph.2006.08.009

17. Chen, X., Margolis, K.J., Gershon, M.D., Schwartz, G.J., Sze, J.Y., 2012. Reduced serotonin reuptake transporter (SERT) function causes insulin resistance and hepatic steatosis independent of food intake. PLoS One. https://doi.org/10.1371/journal.pone.0032511

18. Coppen, A., 1967. The biochemistry of affective disorders. Br. J. Psychiatry. https://doi.org/10.1192/bjp.113.504.1237

19. de Kloet, E.R., Joëls, M., Holsboer, F., 2005. Stress and the brain: from adaptation to disease. Nat. Rev. Neurosci. 6, 463–75. https://doi.org/10.1038/nrn1683

20. Deltheil, T., Guiard, B.P.P., Cerdan, J., David, D.J.J., Tanaka, K.F.F., Repérant, C., Guilloux, J.-P.P., Coudoré, F., Hen, R., Gardier, A.M.M., 2008. Behavioral and serotonergic consequences of decreasing or increasing hippocampus brain-derived neurotrophic factor protein levels in mice. Neuropharmacology 55, 1006–1014. https://doi.org/10.1016/j.neuropharm.2008.08.001

21. Duclot, F., Kabbaj, M., 2015. Epigenetic mechanisms underlying the role of brain-derived neurotrophic factor in depression and response to antidepressants. J. Exp. Biol. 218, 21–31. https://doi.org/10.1242/jeb.107086

22. Duman, R.S., Malberg, J., Nakagawa, S., D’Sa, C., 2000. Neuronal plasticity and survival in mood disorders. Biol. Psychiatry 48, 732–739. https://doi.org/10.1016/S0006-3223(00)00935-5

23. Duman, R.S., Malberg, J., Thome, J., 1999. Neural plasticity to stress and antidepressant treatment. Biol. Psychiatry 46, 1181–1191. https://doi.org/10.1016/S0006-3223(99)00177-8

24. Duman, R.S., Monteggia, L.M., 2006. A Neurotrophic Model for Stress-Related Mood Disorders. Biol. Psychiatry 59, 1116–1127. https://doi.org/10.1016/j.biopsych.2006.02.013

25. Dwivedi, Y., Rizavi, H.S., Conley, R.R., Roberts, R.C., Tamminga, C.A., Pandey, G.N., 2003. Altered Gene Expression of Brain-Derived Neurotrophic Factor and Receptor Tyrosine Kinase B in Postmortem Brain of Suicide Subjects. Arch. Gen. Psychiatry 60, 804. https://doi.org/10.1001/archpsyc.60.8.804

26. Edelmann, E., Leßmann, V., Brigadski, T., 2014. Pre- and postsynaptic twists in BDNF secretion and action in synaptic plasticity. Neuropharmacology 76, 610–627. https://doi.org/10.1016/j.neuropharm.2013.05.043

27. Elbejjani, M., Fuhrer, R., Abrahamowicz, M., Mazoyer, B., Crivello, F., Tzourio, C., Dufouil, C., 2015. Depression, depressive symptoms, and rate of hippocampal atrophy in a longitudinal cohort of older men and women. Psychol. Med. 45, 1931–1944. https://doi.org/10.1017/S0033291714003055

28. Elbejjani, M., Fuhrer, R., Abrahamowicz, M., Mazoyer, B., Crivello, F., Tzourio, C., Dufouil, C., 2014. Hippocampal atrophy and subsequent depressive symptoms in older men and women: Results from a 10-year prospective cohort. Am. J. Epidemiol. 180, 385–393. https://doi.org/10.1093/aje/kwu132

29. Fanelli, G., Serretti, A., 2019. The influence of the serotonin transporter gene 5-HTTLPR polymorphism on suicidal behaviors: a meta-analysis. Prog. Neuro-Psychopharmacology Biol. Psychiatry 88, 375–387. https://doi.org/10.1016/J.PNPBP.2018.08.007

30. Gotlib, I.H., Joormann, J., Minor, K.L., Hallmayer, J., 2008. HPA Axis Reactivity: A Mechanism Underlying the Associations Among 5-HTTLPR, Stress, and Depression. Biol. Psychiatry. https://doi.org/10.1016/j.biopsych.2007.10.008

31. Guidotti, G., Calabrese, F., Auletta, F., Olivier, J., Racagni, G., Homberg, J., Riva, M.A., 2012. Developmental Influence of the Serotonin Transporter on the Expression of Npas4 and GABAergic Markers: Modulation by Antidepressant Treatment. Neuropsychopharmacology 37, 746–758. https://doi.org/10.1038/npp.2011.252

32. Homberg, J.R., De Boer, S.F., Raasø, H.S., Olivier, J.D.A., Verheul, M., Ronken, E., Cools, A.R., Ellenbroek, B.A., Schoffelmeer, A.N.M., Vanderschuren, L.J.M.J., De Vries, T.J., Cuppen, E., 2008. Adaptations in pre- and postsynaptic 5-HT1A receptor function and cocaine supersensitivity in serotonin transporter knockout rats. Psychopharmacology (Berl). 200, 367–380. https://doi.org/10.1007/s00213-008-1212-x

33. Homberg, J.R., Molteni, R., Calabrese, F., Riva, M.A., 2014. The serotonin-BDNF duo: Developmental implications for the vulnerability to psychopathology. Neurosci. Biobehav. Rev. 43, 35–47. https://doi.org/10.1016/j.neubiorev.2014.03.012

34. Jacobs, B.L., van Praag, H., Gage, F.H., 2000. Adult brain neurogenesis and psychiatry: a novel theory of depression. Mol. Psychiatry 5, 262–269. https://doi.org/10.1038/sj.mp.4000712

35. Jansen, F., Heiming, R.S., Lewejohann, L., Touma, C., Palme, R., Schmitt, A., Lesch, K.P., Sachser, N., 2010. Modulation of behavioural profile and stress response by 5-HTT genotype and social experience in adulthood. Behav. Brain Res. https://doi.org/10.1016/j.bbr.2009.09.033

36. Kalueff, A. V., Olivier, J.D.A., Nonkes, L.J.P., Homberg, J.R., 2010. Conserved role for the serotonin transporter gene in rat and mouse neurobehavioral endophenotypes. Neurosci. Biobehav. Rev. 34, 373–386. https://doi.org/10.1016/j.neubiorev.2009.08.003

37. Karpova, N.N., Lindholm, J., Pruunsild, P., Timmusk, T., Castrén, E., 2009. Long-lasting behavioural and molecular alterations induced by early postnatal fluoxetine exposure are restored by chronic fluoxetine treatment in adult mice. Eur. Neuropsychopharmacol. https://doi.org/10.1016/j.euroneuro.2008.09.002

38. Klein Gunnewiek, T.M.M., Homberg, J.R.R., Kozicz, T., 2018. Modulation of glucocorticoids by the serotonin transporter polymorphism: A narrative review. Neurosci. Biobehav. Rev. 92, 338–349. https://doi.org/10.1016/j.neubiorev.2018.06.022

39. Krishnan, V., Nestler, E.J., 2008. The molecular neurobiology of depression. Nature. https://doi.org/10.1038/nature07455

40. Kroeze, Y., Zhou, H., Homberg, J.R., 2012. The genetics of selective serotonin reuptake inhibitors. Pharmacol. Ther. 136, 375–400. https://doi.org/10.1016/J.PHARMTHERA.2012.08.015

41. Lesch, K.P.K.-P., Bengel, D., Heils, A., Sabol, S.Z., Greenberg, B.D., Petri, S., Benjamin, J., Muller, C.R., Hamer, D.H., Murphy, D.L., Müller, C.R., Hamer, D.H., Murphy, D.L., 1996. Association of anxiety-related traits with a polymorphism in the serotonin transporter gene regulatory region. Science (80-.). 274, 1527–1531. https://doi.org/10.1126/science.274.5292.1527

42. Li, Q., Wichems, C., Heils, A., Van De Kar, L.D., Lesch, K.P., Murphy, D.L., 1999. Reduction of 5-hydroxytryptamine (5-HT)(1A)-mediated temperature and neuroendocrine responses and 5-HT(1A) binding sites in 5-HT transporter knockout mice. J. Pharmacol. Exp. Ther.

43. Lira, A., Zhou, M., Castanon, N., Ansorge, M.S., Gordon, J.A., Francis, J.H., Bradley-Moore, M., Lira, J., Underwood, M.D., Arango, V., Kung, H.F., Hofer, M.A., Hen, R., Gingrich, J.A., 2003. Altered depression-related behaviors and functional changes in the dorsal raphe nucleus of serotonin transporter-deficient mice. Biol. Psychiatry 54, 960–971. https://doi.org/10.1016/S0006-3223(03)00696-6

44. Liu, W., Ge, T., Leng, Y., Pan, Z., Fan, J., Yang, W., Cui, R., 2017. The Role of Neural Plasticity in Depression: From Hippocampus to Prefrontal Cortex. Neural Plast. 2017, 1–11. https://doi.org/10.1155/2017/6871089

45. Luoni, A., Hulsken, S., Cazzaniga, G., Racagni, G., Homberg, J.R., Riva, M.A., 2013. Behavioural and neuroplastic properties of chronic lurasidone treatment in serotonin transporter knockout rats. Int. J. Neuropsychopharmacol. https://doi.org/10.1017/S1461145712001332

46. Manfré, G., Clemensson, E.K.H., Kyriakou, E.I., Clemensson, L.E., Van Der Harst, J.E., Homberg, J.R., Nguyen, H.P., 2017. The BACHD rat model of huntington disease shows specific deficits in a test battery of motor function. Front. Behav. Neurosci. https://doi.org/10.3389/fnbeh.2017.00218

47. Mateos-Aparicio, P., Rodríguez-Moreno, A., 2019. The impact of studying brain plasticity. Front. Cell. Neurosci. https://doi.org/10.3389/fncel.2019.00066

48. Matsuda, N., Lu, H., Fukata, Y., Noritake, J., Gao, H., Mukherjee, S., Nemoto, T., Fukata, M., Poo, M. -m. M.M., 2009. Differential Activity-Dependent Secretion of Brain-Derived Neurotrophic Factor from Axon and Dendrite. J. Neurosci. 29, 14185–14198. https://doi.org/10.1523/JNEUROSCI.1863-09.2009

49. Mattson, M.P., Maudsley, S., Martin, B., 2004. BDNF and 5-HT: a dynamic duo in age-related neuronal plasticity and neurodegenerative disorders. Trends Neurosci. 27, 589–594. https://doi.org/10.1016/j.tins.2004.08.001

50. Maynard, K.R., Hill, J.L., Calcaterra, N.E., Palko, M.E., Kardian, A., Paredes, D., Sukumar, M., Adler, B.D., Jimenez, D. V., Schloesser, R.J., Tessarollo, L., Lu, B., Martinowich, K., 2016. Functional Role of BDNF Production from Unique Promoters in Aggression and Serotonin Signaling. Neuropsychopharmacology 41, 1943–1955. https://doi.org/10.1038/npp.2015.349

51. Menzaghi, F., Heinrichs, S.C., Merlo-Pich, E., Tilders, F.J.H., Koob, G.F., 1994. Involvement of hypothalamic corticotropin-releasing factor neurons in behavioral responses to novelty in rats. Neurosci. Lett. 168, 139–142. https://doi.org/10.1016/0304-3940(94)90435-9

52. Miceli, S., Nadif Kasri, N., Joosten, J., Huang, C., Kepser, L., Proville, R., Selten, M.M., van Eijs, F., Azarfar, A., Homberg, J.R., Celikel, T., Schubert, D., 2017. Reduced Inhibition within Layer IV of Sert Knockout Rat Barrel Cortex is Associated with Faster Sensory Integration. Cereb. Cortex 27, 933–949. https://doi.org/10.1093/cercor/bhx016

53. Molteni, R., Cattaneo, A., Calabrese, F., Macchi, F., Olivier, J.D.A., Racagni, G., Ellenbroek, B.A., Gennarelli, M., Riva, M.A., 2010. Reduced function of the serotonin transporter is associated with decreased expression of BDNF in rodents as well as in humans. Neurobiol. Dis. 37, 747–755. https://doi.org/10.1016/j.nbd.2009.12.014

54. Monteggia, L.M., Barrot, M., Powell, C.M., Berton, O., Galanis, V., Gemelli, T., Meuth, S., Nagy, A., Greene, R.W., Nestler, E.J., 2004. Essential role of brain-derived neurotrophic factor in adult hippocampal function. Proc. Natl. Acad. Sci. U. S. A. https://doi.org/10.1073/pnas.0402141101

55. Morris, M.C., Mielock, A., Rao, U., 2017. The SAGE Encyclopedia of Abnormal and Clinical Psychology. https://doi.org/10.4135/9781483365817 NV - 7

56. Murphy, D.L., Fox, M.A., Timpano, K.R., Moya, P.R., Ren-Patterson, R., Andrews, A.M., Holmes, A., Lesch, K.-P.P., Wendland, J.R., 2008. How the serotonin story is being rewritten by new gene-based discoveries principally related to SLC6A4, the serotonin transporter gene, which functions to influence all cellular serotonin systems. Neuropharmacology 55, 932–960. https://doi.org/10.1016/j.neuropharm.2008.08.034

57. Nestler, E.J., Barrot, M., DiLeone, R.J., Eisch, A.J., Gold, S.J., Monteggia, L.M., 2002. Neurobiology of Depression. Neuron 34, 13–25. https://doi.org/10.1016/S0896-6273(02)00653-0

58. O’Hara, R., Schröder, C.M., Mahadevan, R., Schatzberg, A.F., Lindley, S., Fox, S., Weiner, M., Kraemer, H.C., Noda, A., Lin, X., Gray, H.L., Hallmayer, J.F., 2007. Serotonin transporter polymorphism, memory and hippocampal volume in the elderly: Association and interaction with cortisol. Mol. Psychiatry. https://doi.org/10.1038/sj.mp.4001978

59. O’Leary, O.F., Cryan, J.F., O’Leary, O.F., Cryan, J.F., 2014. A ventral view on antidepressant action: Roles for adult hippocampal neurogenesis along the dorsoventral axis. Trends Pharmacol. Sci. 35, 675–687. https://doi.org/10.1016/j.tips.2014.09.011

60. Olivier, J.D.A., Van Der Hart, M.G.C., Van Swelm, R.P.L., Dederen, P.J., Homberg, J.R., Cremers, T., Deen, P.M.T., Cuppen, E., Cools, A.R., Ellenbroek, B.A., 2008. A study in male and female 5-HT transporter knockout rats: An animal model for anxiety and depression disorders. Neuroscience 152, 573–584. https://doi.org/10.1016/J.NEUROSCIENCE.2007.12.032

61. Olivier, J.D.A.D.A., Van Der Hart, M.G.C.G.C., Van Swelm, R.P.L.P.L., Dederen, P.J.J., Homberg, J.R.R., Cremers, T., Deen, P.M.T.M.T., Cuppen, E., Cools, A.R.R., Ellenbroek, B.A.A., 2008. A study in male and female 5-HT transporter knockout rats: An animal model for anxiety and depression disorders. Neuroscience 152, 573–584. https://doi.org/10.1016/j.neuroscience.2007.12.032

62. Pariante, C.M., Lightman, S.L., 2008. The HPA axis in major depression: classical theories and new developments, Trends in Neurosciences. Elsevier Current Trends. https://doi.org/10.1016/j.tins.2008.06.006

63. Park, H., Poo, M.M., 2013. Neurotrophin regulation of neural circuit development and function. Nat. Rev. Neurosci. https://doi.org/10.1038/nrn3379

64. Pawluski, J.L., Kumar, N., Boulle, F., Steinbusch, H.W.M., Machiels, B., Homberg, J.R., Kroeze, Y., van den Hove, D.L.A., Kenis, G., Pawluski, J.L., Homberg, J.R., Machiels, B., Kroeze, Y., Kumar, N., Steinbusch, H.W.M., Kenis, G., van den Hove, D.L.A., 2016. Developmental fluoxetine exposure increases behavioral despair and alters epigenetic regulation of the hippocampal BDNF gene in adult female offspring. Horm. Behav. 80, 47–57. https://doi.org/10.1016/j.yhbeh.2016.01.017

65. Paxinos, G., Watson, C., 2007. The Rat Brain in Stereotaxic Coordinates Sixth Edition. Elsevier Acad. Press.

66. Porsolt, R.D., Anton, G., Blavet, N., Jalfre, M., 1978. Behavioural despair in rats: A new model sensitive to antidepressant treatments. Eur. J. Pharmacol. 47, 379–391. https://doi.org/10.1016/0014-2999(78)90118-8

67. Porsolt, R.D., Le Pichon, M., Jalfre, M., 1977. Depression: a new animal model sensitive to antidepressant treatments. Nature 266, 730–732. https://doi.org/10.1038/266730a0

68. Prut, L., Belzung, C., 2003. The open field as a paradigm to measure the effects of drugs on anxiety-like behaviors: a review. Eur. J. Pharmacol. 463, 3–33. https://doi.org/10.1016/S0014-2999(03)01272-X

69. Quesseveur, G., David, D.J., Gaillard, M.C., Pla, P., Wu, M. V., Nguyen, H.T., Nicolas, V., Auregan, G., David, I., Dranovsky, A., Hantraye, P., Hen, R., Gardier, A.M., Déglon, N., Guiard, B.P., 2013. BDNF overexpression in mouse hippocampal astrocytes promotes local neurogenesis and elicits anxiolytic-like activities. Transl. Psychiatry 3. https://doi.org/10.1038/tp.2013.30

70. Ray, M.T., Weickert, C.S., Wyatt, E., Webster, M.J., 2011. Decreased BDNF, trkB-TK+ and GAD67 mRNA expression in the hippocampus of individuals with schizophrenia and mood disorders. J. Psychiatry Neurosci. https://doi.org/10.1503/jpn.100048

71. Rossi, C., Angelucci, A., Costantin, L., Braschi, C., Mazzantini, M., Babbini, F., Fabbri, M.E., Tessarollo, L., Maffei, L., Berardi, N., Caleo, M., 2006. Brain-derived neurotrophic factor (BDNF) is required for the enhancement of hippocampal neurogenesis following environmental enrichment. Eur. J. Neurosci. 24, 1850–1856. https://doi.org/10.1111/j.1460-9568.2006.05059.x

72. Sakata, K., Duke, S.M., 2014. Lack of BDNF expression through promoter IV disturbs expression of monoamine genes in the frontal cortex and hippocampus. Neuroscience 260, 265–275. https://doi.org/10.1016/j.neuroscience.2013.12.013

73. Sakata, K., Jin, L., Jha, S., 2010. Lack of promoter IV-driven BDNF transcription results in depression-like behavior. Genes, Brain Behav. 9, 712–721. https://doi.org/10.1111/j.1601-183X.2010.00605.x

74. Santos, M.A.O., Bezerra, L.S., Carvalho, A.R.M.R., Brainer-Lima, A.M., 2018. Global hippocampal atrophy in major depressive disorder: a meta-analysis of magnetic resonance imaging studies. Trends Psychiatry Psychother. 40, 369–378. https://doi.org/10.1590/2237-6089-2017-0130

75. Saraiva, C., Praça, C., Ferreira, R., Santos, T., Ferreira, L., Bernardino, L., 2016. Nanoparticle-mediated brain drug delivery: Overcoming blood-brain barrier to treat neurodegenerative diseases. J. Control. Release 235, 34–47. https://doi.org/10.1016/j.jconrel.2016.05.044

76. Schipper, P., Brivio, P., de Leest, D., Madder, L., Asrar, B., Rebuglio, F., Verheij, M.M.M., Kozicz, T., Riva, M.A., Calabrese, F., Henckens, M.J.A.G., Homberg, J.R., 2019. Impaired Fear Extinction Recall in Serotonin Transporter Knockout Rats Is Transiently Alleviated during Adolescence. Brain Sci. 9, 118. https://doi.org/10.3390/brainsci9050118

77. Schipper, P., Nonkes, L.J.P., Karel, P., Kiliaan, A.J., Homberg, J.R., 2011. Serotonin transporter genotype x construction stress interaction in rats. Behav. Brain Res. 223, 169–175. https://doi.org/10.1016/j.bbr.2011.04.037

78. Schulz, P.E., Arora, G., 2015. Depression. Contin. Lifelong Learn. Neurol. https://doi.org/10.1212/01.CON.0000466664.35650.b4

79. Sen, S., Duman, R., Sanacora, G., 2008. Serum Brain-Derived Neurotrophic Factor, Depression, and Antidepressant Medications: Meta-Analyses and Implications. Biol. Psychiatry 64, 527–532. https://doi.org/10.1016/J.BIOPSYCH.2008.05.005

80. Smits, B.M.G., Mudde, J.B., Van De Belt, J., Verheul, M., Olivier, J., Homberg, J., Guryev, V., Cools, A.R., Ellenbroek, B.A., Plasterk, R.H.A., Cuppen, E., 2006. Generation of gene knockouts and mutant models in the laboratory rat by ENU-driven target-selected mutagenesis. Pharmacogenet. Genomics. https://doi.org/10.1097/01.fpc.0000184960.82903.8f

81. Taylor, W.D., McQuoid, D.R., Payne, M.E., Zannas, A.S., MacFall, J.R., Steffens, D.C., 2014. Hippocampus Atrophy and the Longitudinal Course of Late-life Depression. Am. J. Geriatr. Psychiatry 22, 1504–1512. https://doi.org/10.1016/J.JAGP.2013.11.004

82. Tjurmina, O.A., Armando, I., Saavedra, J.M., Goldstein, D.S., Murphy, D.L., 2002. Exaggerated adrenomedullary response to immobilization in mice with targeted disruption of the serotonin transporter gene. Endocrinology. https://doi.org/10.1210/en.2002-220416

83. Van den Hove, D.L.A., Kenis, G., Brass, A., Opstelten, R., Rutten, B.P.F., Bruschettini, M., Blanco, C.E., Lesch, K.P., Steinbusch, H.W.M., Prickaerts, J., 2013. Vulnerability versus resilience to prenatal stress in male and female rats; Implications from gene expression profiles in the hippocampus and frontal cortex. Eur. Neuropsychopharmacol. 23, 1226–1246. https://doi.org/10.1016/J.EURONEURO.2012.09.011

84. van der Doelen, R.H.A., Deschamps, W., D’Annibale, C., Peeters, D., Wevers, R.A., Zelena, D., Homberg, J.R., Kozicz, T., 2014. Early life adversity and serotonin transporter gene variation interact at the level of the adrenal gland to affect the adult hypothalamopituitary-adrenal axis. Transl. Psychiatry 4, e409–e409. https://doi.org/10.1038/tp.2014.57

85. Vertes, R.P., 2004. Differential Projections of the Infralimbic and Prelimbic Cortex in the Rat. Synapse 51, 32–58. https://doi.org/10.1002/syn.10279

86. Verwer, R.W.H., Meijer, R.J., Van Uum, H.F.M., Witter, M.P., 1997. Collateral projections from the rat hippocampal formation to the lateral and medial prefrontal cortex. Hippocampus. https://doi.org/10.1002/(SICI)1098-1063(1997)7:4<397::AID-HIPO5>3.0.CO;2-G

87. Vidal-Gonzalez, I., Vidal-Gonzalez, B., Rauch, S.L., Quirk, G.J., 2006. Microstimulation reveals opposing influences of prelimbic and infralimbic cortex on the expression of conditioned fear. Learn. Mem. https://doi.org/10.1101/lm.306106

